# A novel approach for assessing hypoperfusion in stroke using spatial independent component analysis of resting-state fMRI

**DOI:** 10.1101/2020.07.17.208058

**Authors:** Jiun-Yiing Hu, Evgeniya Kirilina, Till Nierhaus, Dr. rer. medic. Dipl.-Ing.(FH), Smadar Ovadia-Caro, Michelle Livne, Kersten Villringer, Daniel Margulies, Jochen B. Fiebach, Arno Villringer, Ahmed A. Khalil

**Affiliations:** Department of Internal Medicine, University of Maryland School of Medicine, Baltimore, USA; Department of Neurophysics, Max-Planck-Institute for Human Cognitive and Brain Sciences, Leipzig, Germany; Neurocomputation and Neuroimaging Unit, Center for Cognitive Neuroscience Berlin (CCNB), Department of Education and Psychology, Freie Universität Berlin, Germany; Department of Neurology, Max-Planck-Institute for Human Cognitive and Brain Sciences, Leipzig, Germany; Department of Cognitive Sciences, University of Haifa, Israel; Charité-Universitätsmedizin Berlin, Center for Stroke Research Berlin, Germany; Centre National de la Recherche Scientifique (CNRS) UMR 7225, Frontlab, Institut du Cerveau et de la Moelle Épinière, Paris, France; Berlin School of Mind and Brain, Humboldt Universität zu Berlin, Berlin, Germany; Berlin Institute of Health (BIH), Berlin, Germany

**Keywords:** blood oxygenation level dependent signal, perfusion, resting-state functional magnetic resonance imaging, spatial independent component analysis, stroke

## Abstract

Individualized treatment of acute stroke depends on the timely detection of ischemia and potentially salvageable tissue in the brain. Using functional MRI (fMRI), it is possible to characterize cerebral blood flow from blood-oxygen-level-dependent (BOLD) signals without the administration of exogenous contrast agents. In this study, we applied spatial independent component analysis to resting-state fMRI data of 37 stroke patients scanned within 24 hours of symptom onset, 17 of whom received follow-up scans the next day. Our analysis revealed “Hypoperfusion spatially-Independent Components” (HICs) whose spatial patterns of BOLD signal resembled regions of delayed perfusion depicted by dynamic susceptibility contrast MRI. These HICs were detected even in the presence of excessive patient motion, and disappeared following successful tissue reperfusion. The unique spatial and temporal features of HICs allowed them to be distinguished with high accuracy from other components in a user-independent manner (AUC = 0.95, accuracy = 0.96, sensitivity = 1.00, specificity = 0.96). Our study therefore presents a new, non-invasive method for assessing blood flow in acute stroke that minimizes interpretative subjectivity and is robust to severe patient motion.

## 1 Introduction

Imaging blood flow in acute ischemic stroke is important in directing treatment decisions. In clinical practice, MRI-based blood flow imaging is typically performed using dynamic susceptibility contrast MRI (DSC-MRI). This method is well-established and correlates well with the in-vivo perfusion imaging gold-standard, ^15^O-water positron emission tomography (PET) (Zaro-Weber et al., 2010, 2016). However, it requires the use of intravenously administered contrast agents, which are contraindicated in patients with kidney impairment (Khawaja et al., 2015) and can accumulate in tissues with repeated use (Gulani et al., 2017).

BOLD functional MRI has been proposed as an alternative to DSC-MRI. At rest, BOLD signal oscillations reflect a combination of different signal sources, including the hemodynamic response to spontaneous neural activity (neurovascular coupling), peripheral physiology (respiratory and cardiac function), fluctuations in blood carbon dioxide (Golestani et al., 2015; Wise et al., 2004), and head motion (T. T. Liu, 2016). Disturbed cerebral blood flow is associated with changes in BOLD signal fluctuations in multiple ways. One of the best studied perfusion-related changes in the BOLD signal is BOLD delay (Amemiya et al., 2014; Q. Chen et al., 2018; Christen et al., 2015; Khalil et al., 2017, 2020; Lv et al., 2013, 2018; Ni et al., 2017; Siegel et al., 2016; J. Wu et al., 2017), which are relative phase lags in low frequency oscillations (LFOs) of the BOLD signal originating from outside the brain (systemic LFOs) in underperfused brain regions (Tong et al., 2019). Disturbed perfusion is known to affect other BOLD signal features, including a shift to lower LFO frequencies (Y. Liu et al., 2007; Tsai et al., 2014; Yao et al., 2012) and a reduction in LFO amplitude (Wang et al., 2008). These findings suggest that regions of disturbed perfusion may have a unique set of BOLD signal characteristics that distinguish them from normally perfused tissue. Using this BOLD “signature” to rapidly and reliably identify abnormally perfused tissue would be of great diagnostic and prognostic utility.

Spatial independent component analysis (spatial ICA) is a data-driven method for decomposing complex signals into statistically-independent subcomponents (Beckmann & Smith, 2004). In fMRI, it has been used to remove non-neuronal signal contributions such as head motion, large vessel pulsations, and scanner instabilities (Thomas et al., 2002). Using spatial ICA, researchers can isolate functionally distinct resting-state networks (RSNs) such as visual cortical areas and the sensory-motor cortex (Beckmann et al., 2005).

Considering spatial ICA’s ability to separate statistically-independent sources of variance in the BOLD signal, we investigated whether spatial ICA would be able to detect hypoperfusion-related changes in the BOLD signal in acute stroke patients. Specifically, we aimed 1) to identify spatial ICA components corresponding to tissue hypoperfusion, 2) to characterize the temporal and spatial properties of these “Hypoperfusion spatially-Independent Components” (HICs), and 3) to determine whether these properties allow the automated identification of HICs.

## 2 Materials and Methods

### 2.1 Patients

This study analyzes data from a subset of ischemic stroke patients from 1000Plus (Hotter et al., 2009) (clinicaltrials.gov identifier: NCT00715533) and LOBI-BBB (clinicaltrials.gov identifier: NCT02077582), two prospective cohort imaging studies conducted at Charité Universitätsmedizin Campus Benjamin Franklin (Berlin, Germany) between January 2009 and March 2016.

The patients included in this study had a clinical and radiological diagnosis of acute supratentorial ischemic stroke, and had received both dynamic susceptibility contrast magnetic resonance imaging (DSC-MRI) and resting-state functional MRI (rs-fMRI) scans within 24 hours of symptom onset as part of a standard stroke imaging protocol (Hotter et al., 2009). All patients showed perfusion deficits on their time-to-maximum (Tmax) maps at baseline. Where available, follow-up scans from the same patients, acquired approximately 24 hours after the baseline scans, were also included in the analysis. Exclusion criteria were any contraindications to undergoing an MRI or receiving a gadolinium contrast agent.

Thirty-seven acute ischemic stroke patients who met our inclusion criteria were included in this study, fourteen of whom had follow-up scans. A total of 51 scans were thus included in data analysis. The baseline scans of twenty-two of these patients were previously analyzed in a comparison between time shift analysis of rs-fMRI data (BOLD delay) and DSC-MRI (Khalil et al., 2017) and twelve patients (and their follow-up scans) from the current study were previously investigated in a study of the longitudinal evolution of perfusion deficits detected using BOLD delay (Khalil et al., 2020).

### 2.2 Ethics statement

Patients provided written informed consent prior to participation, and all procedures were approved by the local ethics committee (EA4/026/08 for the 1000plus study and EA1/200/13 for LOBI-BBB) and were performed according to the Declaration of Helsinki.

### 2.3 Imaging

#### 2.3.1 Acquisition

All imaging was performed on a Siemens (Erlangen, Germany) Tim Trio 3 Tesla MRI scanner. The sequence parameters were as follows, rs-fMRI; repetition time (TR) = 2300 ms, echo time (TE) = 30 ms, flip angle (FA) = 90**°**, matrix = 64 x 64, voxel dimensions = 3 x 3 x 3 mm^3^, 1 mm slice gap, 33 slices and 150 volumes (acquisition time=5 min. and 50 s). The rs-fMRI scan was performed prior to the administration of the contrast agent and patients were requested to relax, lie still, and close their eyes for the duration of the scan.

DSC-MRI was performed after injection of a bolus of 5ml Gadovist ® (Gadobutrol, 1M, Bayer Schering Pharma AG, Berlin, Germany) followed by a saline flush at a flow rate of 5ml/s. The sequence parameters were: TR = 1390 ms, TE = 29 ms, FA = 60, matrix = 128 x 128, voxel dimensions = 1.8 x 1.8 x 5 mm, 0.5 mm slice gap, 21 slices and 80 volumes (acquisition time = 1 min and 58 s).

Other imaging included diffusion-weighted imaging (DWI), time-of-flight MR angiography, and fluid-attenuated inversion recovery (FLAIR) scans, as described previously (Hotter et al., 2009).

#### 2.3.2 Dynamic susceptibility contrast MRI

Maps of time-to-maximum of the tissue residue function (Tmax) were generated from the DSC-MRI data using Stroketool (version 2.8; Digital Image Solutions – H.J. Wittsack) after deconvolution of the concentration-time curve through block-circulant singular value decomposition (O. Wu et al., 2003). The arterial input function (AIF) was selected from 5-10 voxels in the distal branches of the middle cerebral artery on the hemisphere contralateral to the acute infarct (Ebinger et al., 2010). Artifacts in the cerebrospinal fluid (CSF) were automatically removed from the Tmax maps using CSF masks derived from T2-weighted B0 images.

#### 2.3.3 Resting-state functional MRI

##### 2.3.3.1 Preprocessing

Preprocessing of the resting-state data was performed using tools from Analysis of Functional NeuroImages (AFNI) (Cox, 1996) and the FMRIB Software Library (FSL). The first four volumes of each time series were discarded for signal equilibration. After correcting for slice-timing effects, the remaining volumes were skull-stripped, realigned to the mean functional image of each individual, and linear and quadratic trends were removed. Images were spatially smoothed (Gaussian kernel of full-width-at-half-maximum = 6 mm), but left temporally unfiltered to include the full range of signal frequencies for spatial ICA.

##### 2.3.3.2 ICA decomposition

Spatial ICA is a data-driven approach for multivariate data analysis and representation. It was implemented in this study using FSL’s Multivariate Exploratory Linear Optimized

Decomposition into Independent Components - MELODIC (Beckmann & Smith, 2004). Spatial ICA models four-dimensional fMRI data as a linear combination of unknown source signals (referred to as “independent components”), each described by a spatial map and a mean time course across all voxels in the spatial map. It dissects out the different source signals by assuming their mutual, statistical independence and non-Gaussianity (Beckmann & Smith, 2004). In this study, spatial ICA was implemented using MELODIC’s default Bayesian dimensionality estimation, which estimates the number of source signals within the data automatically (Beckmann & Smith, 2004). This means that the number of independent components outputted by MELODIC can differ between patients and scanning sessions in the study.

##### 2.3.3.3 Manual classification of independent components

For a subset of the data (20 out of 37 baseline scans; the **FIX training dataset**, see below), two raters (A.A.K and J-Y.H) together (i.e. not independently) manually classified the independent components from MELODIC into the following classes: resting-state networks (RSNs), likely hypoperfusion independent components (HICs), noise (including head motion and scanner noise), or “unknown” if the independent component could not be clearly identified. Manual classification of the RSNs, noise, and unknown components was carried out according to published guidelines (Kelly et al., 2010) by inspecting the thresholded spatial maps (|Z| > 2.33), the power spectrum, and the time courses. HICs were identified as unilateral components that were present in, and largely restricted to, the vascular territory in which the patient’s infarct was found (guided by the DWI).

A total of 896 independent components were generated using MELODIC from the subjects in the FIX training dataset. The median number of independent components per subject was 45 (IQR = 43 – 47) (Supplementary Figure 1). The mean (± SD) contribution of each component class (% out of the total number of components) for subjects in the FIX-training dataset are as follows: noise (79.8% ± 9.4%), resting-state networks (9.2% ± 5.9%), and likely hypoperfusion independent component(s) (HIC; 2.6% ± 1.5%). “Other” components that could not be unambiguously classified into any of the other three categories formed 8.4% ± 4.8% of the training dataset. Of the 20 subjects in the FIX training dataset, 14 had a single HIC, 3 had two HICs, 1 had three HICs, and 2 had no HICs. Examples of our classifications are provided in Supplementary Figure 2.

##### 2.3.3.4 Data denoising

To reduce the number of components by removing non-signal independent components, we used FMRIB’s ICA-based X-noiseifier (FIX), a machine-learning algorithm that automatically classifies signal and artifact components in rs-fMRI data using 186 spatial and temporal features (Salimi-Khorshidi et al., 2014). Minor modifications were made to FIX to accommodate our data: we substituted T1-weighted magnetization-prepared rapid gradient-echo (MPRAGE) scans used in calculations of FIX features with B0 images (T2-weighted spin-echo EPI), as MPRAGEs are not routinely acquired as part of our acute stroke MR imaging protocol (Hotter et al., 2009). Tissue segmentation maps needed for these features were generated from these images using FSL’s FMRIB Automated Segmentation Tool (FAST). All segmentations were visually assessed for quality.

FIX was trained on the FIX training dataset. The accuracy of FIX for separating signal from noise in the FIX training dataset was investigated using leave-one-out cross-validation. A threshold of 20 was selected for the classification of independent components in our training data, as this was the highest threshold at which no HICs were incorrectly classified by FIX as noise (see Supplementary Table 1). At this threshold, FIX achieved a mean true positive rate (TPR; percent of true signal independent components correctly classified) of 90.9%, and a mean true negative rate (TNR; percent of true noise independent components correctly classified) of 84.6%. The output of FIX denoising was a “shortlist” of components classified by FIX as being “non-noise”. After training, FIX was applied to the full dataset (n = 2337 independent components from 51 scans). FIX reduced the number of independent components to a list of likely-signal independent components by 77.4% (total, n = 528 independent components from 51 scans, see Supplementary Figure 3).

##### 2.3.3.5 Time shift analysis

For a visual comparison between HICs and a more established measure of perfusion derived from rs-fMRI, we performed time shift analysis (TSA) of the preprocessed rs-fMRI data (Amemiya et al., 2014; Christen et al., 2015; Khalil et al., 2017, 2020; Lv et al., 2013). TSA maps were generated by assigning each voxel the value of the time shift (ranging between −20 to +20 seconds) that achieves maximum cross-correlation between the voxel’s time series and a recursively refined regressor derived from the global mean time series after univariate interpolation of the time series. A long tracking range was applied because time shift delays in acute stroke patients are very prolonged (Khalil et al., 2017, 2020; Tanrıtanır et al., 2020) TSA was performed using a set of Python tools (https://github.com/bbfrederick/rapidtide)(Frederick, 2016).

#### 2.3.4 Image registration

All images (including Tmax maps, resting-state data, and spatial ICA maps) were registered to a custom echo planar imaging (EPI) template in MNI 152 standard space using a rigid body spatial transformation (3 rotations and 3 translations, implemented using FSL FLIRT) for further processing. This template was derived from the EPI scans of 103 stroke patients and details of how it was generated can be found in the Supplementary Materials of (Khalil et al., 2017). This template was used instead of standard templates, such as the MNI template, to account for differences in brain and ventricular system size between the study’s population and younger, healthy individuals.

### 2.4 Feature extraction

We extracted a set of relevant spatial and temporal features from the shortlisted independent components (i.e. after data denoising by FIX) to examine how HICs may differ from other signal components. The following features were extracted:

1. *Temporal delay to a reference signal:* Temporally delayed BOLD signals, calculated using time shift analysis (Lv et al., 2013), have been found to correspond to hypoperfused brain regions. We thus explored whether the temporal delay between the time courses of each independent component and a reference time course was different between HICs and independent components representing normally perfused tissue. We used two different reference time courses: the whole brain (global) signal (Amemiya et al., 2014; Lv et al., 2013) and the signal from within the major venous sinuses (Aso et al., 2017; Christen et al., 2015; Khalil et al., 2017), the latter of which was automatically extracted using a venous sinus template in MNI space (details described in the Supplementary Materials of (Khalil et al., 2017)). Positive delay values indicate the component’s BOLD signal time course followed the reference time course, while negative delay values indicate the component’s BOLD signal time course preceded the reference time course.
2. *Power in frequency bands:* Ischemia shifts BOLD signal oscillations towards lower frequencies (Y. Liu et al., 2007; Tsai et al., 2014; Yao et al., 2012). For each independent component, we calculated the percentage of total spectral power in six different frequency ranges (0 – 0.01 Hz, 0.01 – 0.025 Hz, 0.025 – 0.05 Hz, 0.05 – 0.1 Hz, 0.1 – 0.15 Hz, 0.15 – 0.20 Hz) after fast Fourier transform of the time courses (Salimi-Khorshidi et al., 2014).
3. *Percent restriction to a single vascular territory:* Ischemic stroke-related hypoperfusion is most commonly restricted to a vascular territory supplied by a single major artery. This makes the percent restriction of an independent component’s spatial map to a single vascular territory a potential feature of interest. We thus calculated the percentage of each independent component’s volume that exists within a single vascular territory. For this analysis, only independent components with >50% of their volume within a single vascular territory were considered. The vascular territories were defined according to a custom vascular territory atlas in MNI template space, described in further detail in the Supplementary Materials of (Khalil et al., 2017).
4. *Mean Tmax delay:* To compare the results of the spatial ICA with the level of perfusion revealed by DSC-MRI in the area covered by each independent component, we calculated the mean Tmax delay of all voxels in the Tmax map overlapped by the independent component’s spatial map.

### 2.5 Automated identification of HICs

#### 2.5.1 Model training

We used a Generalized Linear Model (GLM) on the extracted features to estimate the probability that a given independent component is a HIC. To mitigate multicollinearity effects we applied elastic net regularization to our GLM (Zou & Hastie, 2005).

Disregarding components classified as noise, HICs consisted approximately 10% of the components per dataset. To address this problem of extreme class imbalance (Krawczyk, 2016), which can bias GLM results, we first sub-sampled our dataset to contain 50% HICs (number of components = 57), and 50% non-HICs (number of components = 57). From this **model dataset**, which undersamples the majority (non-HIC) class, we randomly selected 80% of the independent components to be the **training dataset** (containing 43 HICs and 48 non-HICs).

For the model training, components were classified as HICs if their model probabilities were above a probability threshold determined by Youden’s index, which gives equal weight to sensitivity and specificity. The shrinkage parameter lambda was estimated using leave-one-out cross-validation on the training dataset.

#### 2.5.2 Model testing

The **test dataset** contained all the HICs (number of components = 14) and non-HICs (number of components = 414) remaining after selection of the training dataset. From this, we randomly subsampled 5 HICs and a set of non-HICs according to the mean ratio between HICs and non-HICs in individual MELODIC outputs in the FIX shortlist (about 1:10). This random selection was repeated 50 times and the model’s performance is reported as the median of these iterations (see below for metrics of model performance).

### 2.6 Statistical analysis

Radar plots were created using the “radarchart” function from the *fmsb* R package (Nakazawa, 2018) to visualize feature values. Raincloud plots, which combine dot plots, box plots, and violin plots, are used to visualize the distribution of continuous variables (Allen et al., 2019).

The elastic net regularized GLM used in this study was implemented using the “cv.glmnet” function from the *glmnet* R package (Friedman et al., 2010).

Model performance is reported using the following metrics:

- Area-under-the-curve of the receiver operating ROC curve (AUC): The probability that the model will assign a higher probability of being a HIC to a randomly chosen HIC than to a randomly chosen non-HIC. ROC curves were visualized and their AUCs were calculated using the *pROC* R package (Robin et al., 2011)
- Sensitivity: The true positive rate, or the proportion of HICs correctly classified by the model as being HICs
- Specificity: The true negative rate, or the proportion of non-HICs correctly classified by the model as being non-HICs
- Balanced accuracy: (Sensitivity + specificity) / 2. Balanced accuracy was chosen instead of regular accuracy due to the unbalanced nature of our test dataset (Brodersen et al., 2010).
- Cohen’s kappa: The agreement between the model’s classification and the “true”/”reference” classification, beyond that expected due to random chance. This was calculated using the *irr* R package (Gamer et al., 2019).

The GLM coefficients were converted into Odds ratios (Odds ratio = e^coefficient^). Since elastic net, like all penalized regression models, provides biased estimators, no meaningful standard errors can be calculated (Kyung et al., 2010) and therefore p-values are not reported in this analysis. Instead, the model produces penalized coefficients through bootstrapped cross-validation. Variables whose contribution to the outcome is negligible have penalized coefficients of zero (i.e. Odds ratios of 1).

### 2.7 Data/code availability

The data and code for the training and testing of the model can be found at https://github.com/ahmedaak/spatial_ICA_stroke.

## 3 Results

### 3.1 Patients

The study sample consisted of 17 women and 20 men (mean age = 71 years, SD = 14 years). Their median NIHSS was 7 (IQR 3 - 14) and the median time from symptom onset to MRI was 7 hours (IQR 2 - 16 h). Intravenous thrombolysis was administered to 19 (51%) patients. Fourteen patients received follow-up scans on the second day following stroke onset, resulting in a total of 51 scans. Tissue reperfusion was successful in 3 of the 14 patients with follow-up scans, as demonstrated by the complete resolution of time-to-maximum (Tmax) lesions observed on their baseline scans.

Head motion was quantified using framewise displacement (FD), which is the sum of the absolute value of individual subjects’ six translational and rotational realignment parameters (Power et al., 2012). Twelve out of the 51 rsfMRI scans exhibited severe head motion, defined according to previous studies on the use of rsfMRI for assessing cerebral hemodynamics as a mean FD of >0.4 mm across the scan or a maximum FD of >3 mm (Khalil et al., 2017, 2020). Median FD (mean across all scan volumes) in the sample was 0.23 mm (IQR: 0.15 mm) for rs-fMRI, and 0.29 mm (IQR: 0.19 – 0.47 mm) for DSC-MRI. Median FD (maximum across all scan volumes) were 1.28 mm (IQR: 0.61 – 2.86 mm) and 0.82 (IQR: 0.49 – 1.89 mm) for rs-fMRI and DSC-MRI scans, respectively. The distribution of mean and maximum FD for the rs-fMRI scans is shown in Supplementary Figure 4.

### 3.2 Hypoperfusion independent components reflect tissue hypoperfusion

Of the 37 baseline scans, 34 showed at least one HIC (6 scans showed 2 HICs, 2 scans showed 3 HICs). Figure 1 shows the DWI, Tmax, and HIC of three example patients. Maps for the other 31 patients can be found in Supplementary Figure 5. Together, these figures reveal the striking spatial similarities between HICs and perfusion deficits on Tmax maps.

**Figure 1.**
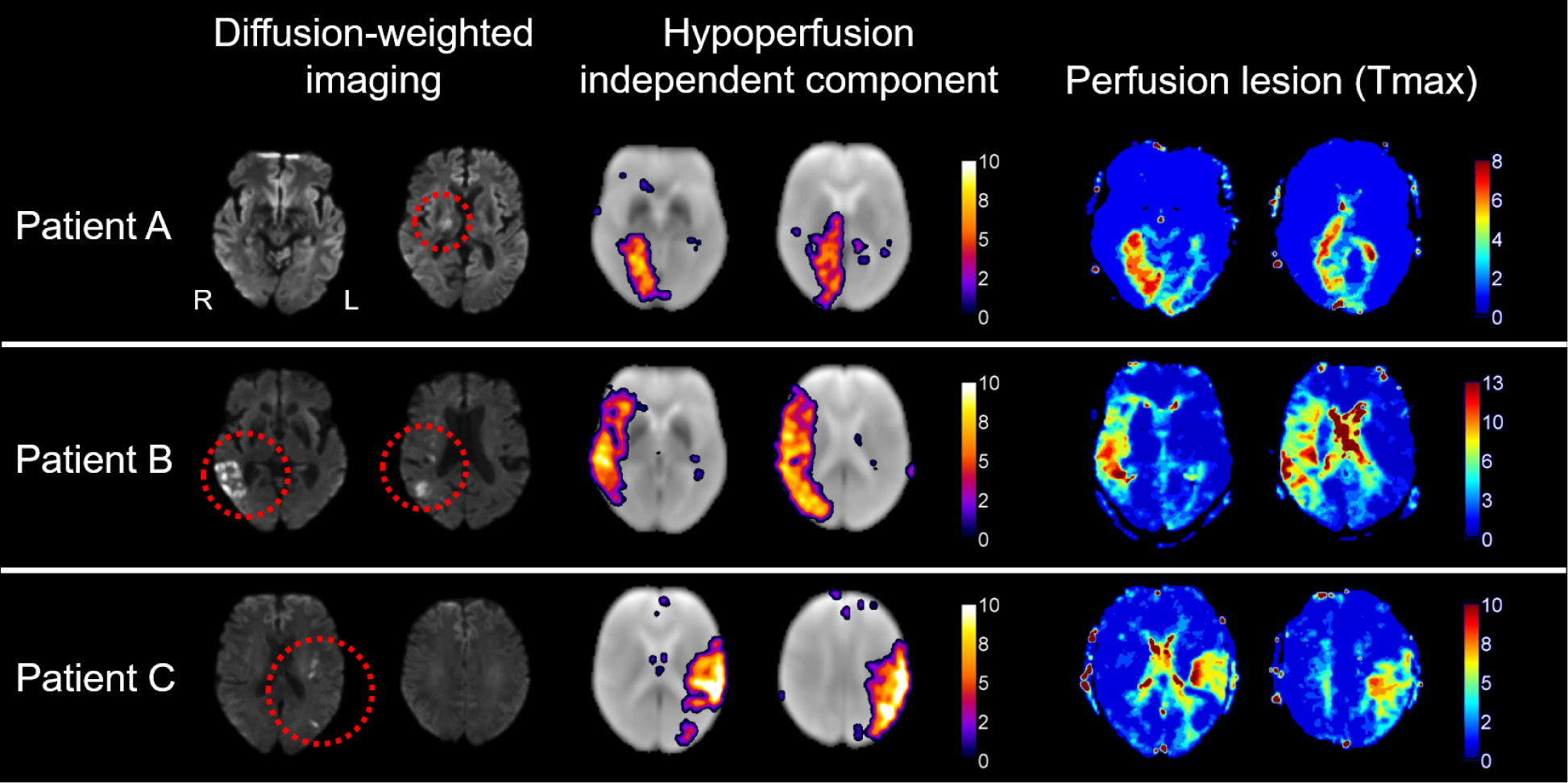
Spatial independent component analysis (spatial ICA) of resting-state fMRI detects post-stroke perfusion deficits in the form of hypoperfusion independent components (HICs). The figure shows (from left to right) diffusion-weighted imaging (visualization of infarcted tissue), hypoperfusion independent components (HICs), and time-to-maximum of the residual curve from dynamic susceptibility contrast MRI (Tmax; reflecting perfusion), from three representative acute stroke patients with a right-sided posterior cerebral artery infarct (patient A), right-sided middle cerebral artery infarct (patient B), and left-sided middle cerebral artery infarct (patient C). A corresponding perfusion deficit is seen in the affected vascular territory of each patient on the Tmax maps. In each case, spatial ICA reveals a HIC with striking spatial resemblance to the perfusion deficit visible on the Tmax maps obtained using a contrast agent.

In addition, 10 out of the 14 follow-up scans showed HICs. Figure 2 illustrates the relationship between the spatial ICA results and vessel status (recanalization or persistent vessel occlusion) in two patients with follow-up scans. In patient A, we observe the resolution of HICs following vessel recanalization and tissue reperfusion, while in patient B, HICs persist with continued vessel occlusion and tissue hypoperfusion. This finding suggests that HICs dynamically respond to changes in vessel status and perfusion dynamics over time.

**Figure 2.**
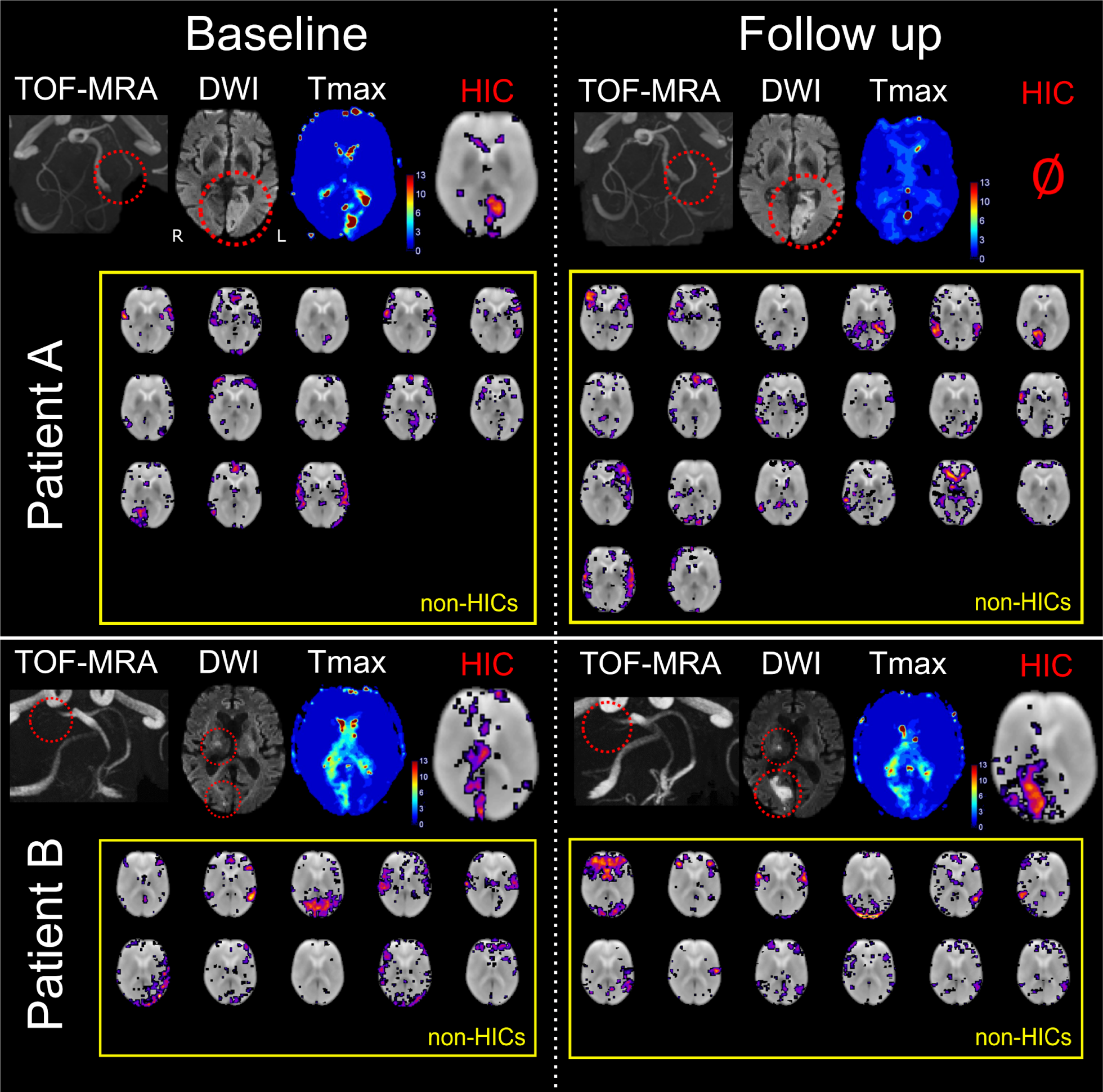
Perfusion assessed using spatial independent component analysis (spatial ICA) evolves with vessel status. Baseline images were obtained within 24 hours of symptom onset, while follow-up images were acquired the next day. Patient A had a left-sided acute posterior cerebral artery (PCA) infarct due to occlusion of the P2 segment of the PCA, seen on time-of-flight magnetic resonance angiography (TOF-MRA) and diffusion weighted imaging (DWI). A perfusion deficit as seen on a contrast-enhanced Tmax map and a corresponding hypoperfusion independent component (HIC) were observed among 13 other independent components (after automatic denoising) at baseline. Following recanalization of the PCA, there is a normalization of the perfusion deficit in the PCA vascular territory on the contrast-enhanced Tmax map. Spatial ICA on follow-up imaging generated 20 independent components after automatic denoising, mostly corresponding to resting-state networks, and no HICs. Patient B had a right-sided acute PCA infarct due to occlusion of the P1 segment. A perfusion deficit and corresponding HIC are seen at baseline. Spatial ICA also produced 10 other independent components that mostly corresponded to resting-state networks (RSNs). Unlike in patient A, patient B’s PCA remained occluded at follow-up. The perfusion deficit in the posterior cerebral artery territory persisted on the Tmax map, and a HIC was observed along with RSN independent components. This demonstrates that HICs follow longitudinal perfusion dynamics corresponding to clinically-relevant events such as changes in vessel status.

HICs were identified in 10 out of 12 subjects with high head motion during their rs-fMRI scans, as shown in an example patient in Figure 3. The images and head motion traces of all 12 subjects with high head motion can be interactively viewed here: https://doi.org/10.6084/m9.figshare.13676779. Overall, there were 4 false negative cases, in which no HIC was visible despite the presence of a perfusion deficit on Tmax maps (these cases can be interactively viewed here https://doi.org/10.6084/m9.figshare.13664384). In two of the false negative cases, extremely severe motion was present in the rs-fMRI data, exceeding 20 mm maximum framewise displacement (see Supplementary Figure 4). In the other two false negative cases, motion during the rs-fMRI scan was much lower (mean FD = 0.25 and 0.14 mm, max FD = 0.56 and 0.44 mm).

**Figure 3.**
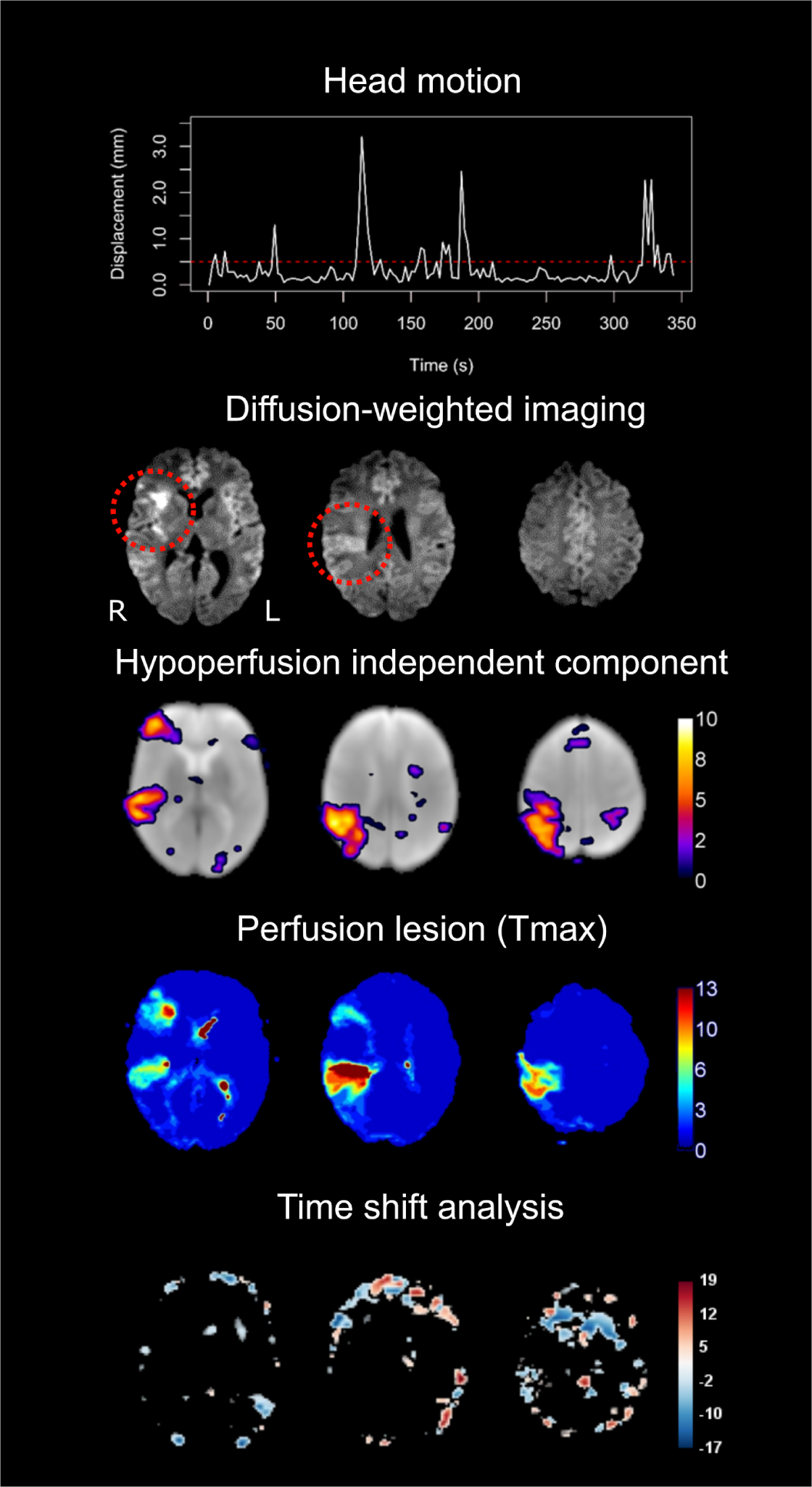
Hypoperfusion independent components (HICs) are detected using spatial ICA despite high head motion. In this patient with a right-sided middle cerebral artery infarct (red circles on the diffusion-weighted image), the top row demonstrates the framewise displacement of the patient’s head over the duration of the resting-state functional MRI scan (maximum motion = 32 mm, mean motion = 0.35 mm). Despite the high motion, a HIC was detected that corresponded to the patient’s perfusion deficit as detected by contrast-enhanced Tmax maps on dynamic susceptibility contrast MRI. No perfusion deficit is seen on the time shift analysis map, derived from the same scan as the HIC. Of note, this patient has two distinct perfusion deficits within the same vascular territory (right middle cerebral artery)—one frontal and one temporoparietal—both of which are reflected in the HIC.

HICs identified multiple perfusion deficits located in different vascular territories within the same patient (Figure 4). The fact that multiple, spatially remote areas of hypoperfusion within an individual patient are captured in a single independent component suggests that these areas share a BOLD signal signature that sets them apart from other components.

**Figure 4.**
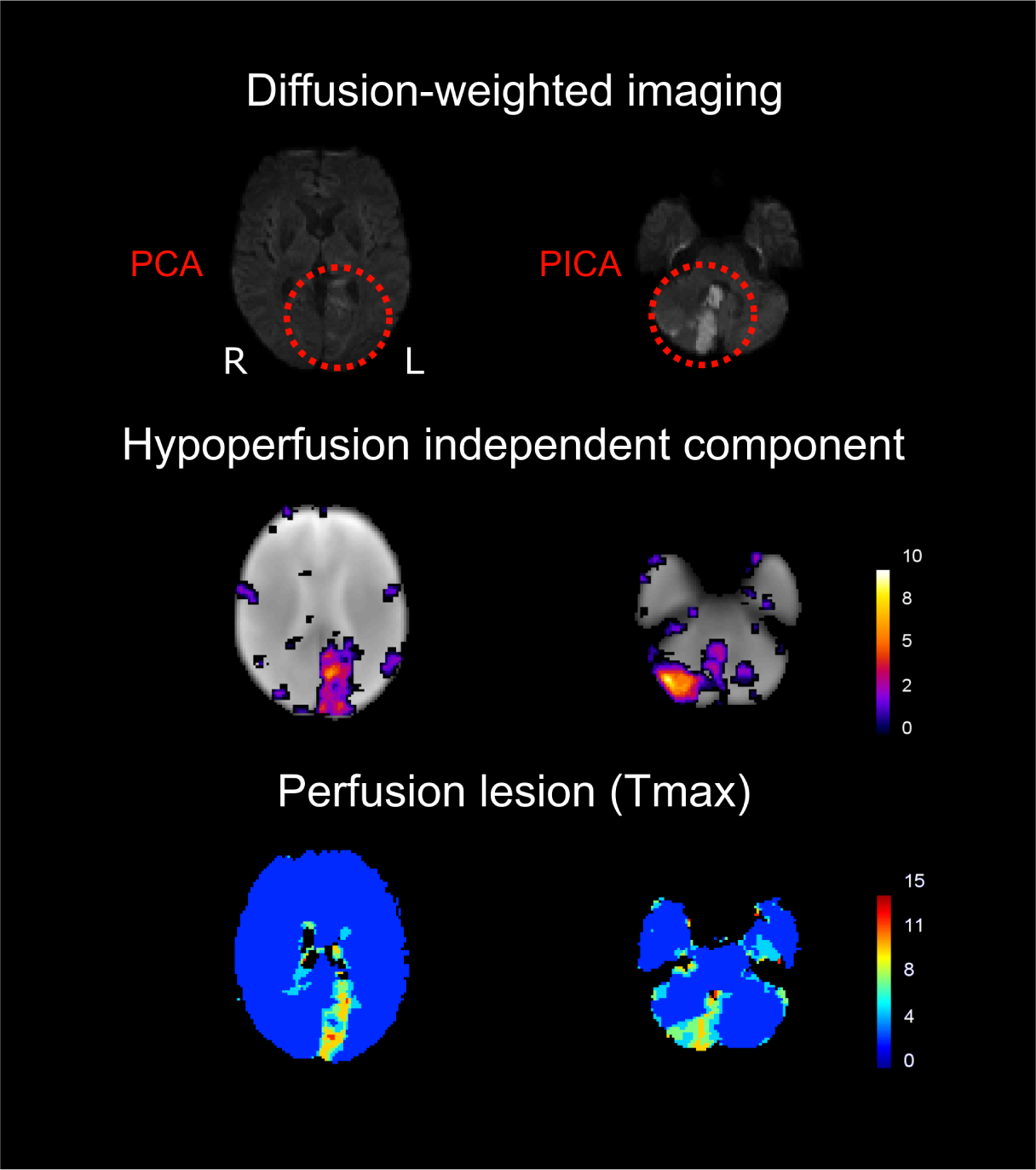
Hypoperfusion independent components (HICs) detect multiple perfusion deficits affecting different vascular territories in the same patient. This patient has multiple infarcts (with corresponding perfusion deficits on the contrast-enhanced Tmax maps - bottom row) - in the left posterior cerebral artery (PCA) territory (right) and in the right posterior inferior cerebellar artery (PICA) territory (left). Spatial independent component analysis reveals a single HIC (middle row) that corresponds to both perfusion deficits in this patient. This implies that these areas of hypoperfusion, although spatially distanced from each other, share a common BOLD signal signature. Note that, in the left column, the DWI slice showing the PCA infarct does not correspond to the same slice on the Tmax and HIC images, as the infarct is not visible on the DWI at the level shown on the Tmax and HIC images.

### 3.2 HICs show unique characteristics

Table 1 shows the feature values (mean across patients) extracted from the independent components.

**Table 1.**
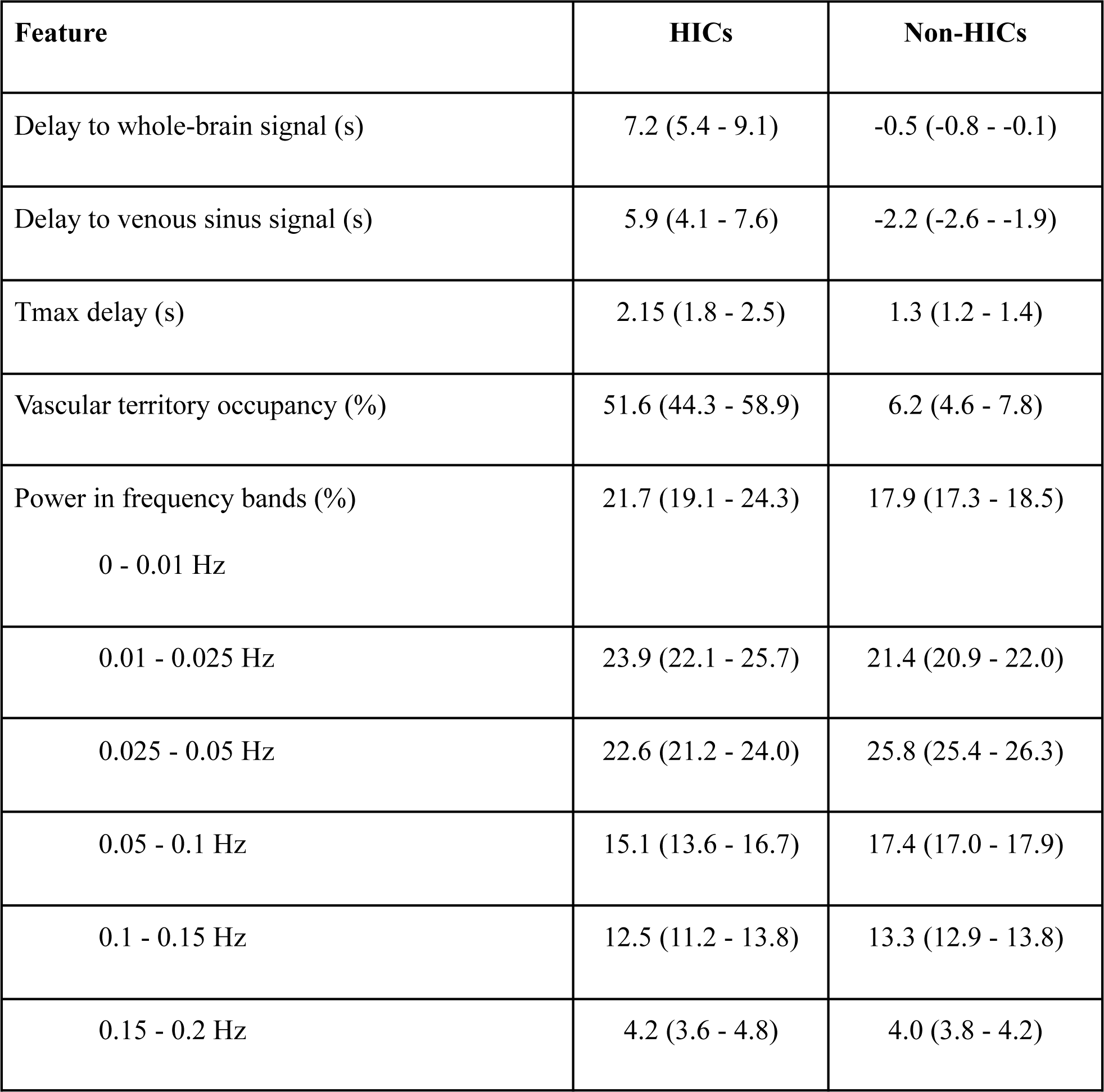
Mean across patients and 95% confidence interval of the feature values (non-normalized) for HICs and non-HICs

For each patient, min-max normalization was applied to the values of the features extracted from each of the independent components using the following equation:

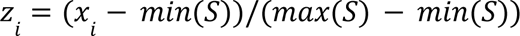

where *z* is the normalized feature value, *x* is the original feature value, *i* refers to the independent component in question, and *S* refers to the feature values of the entire set of independent components from a single patient.

The means of the normalized feature values across patients are shown on the radar plot in Figure 5. HICs have a unique feature signature compared to non-HICs, characterized by delayed low frequency oscillations compared to the whole-brain and venous sinus references, greater restriction to a single vascular territory, greater overlap with areas of hypoperfusion (as assessed using Tmax), and more power in the lowest frequency bands (0 - 0.025 Hz).

**Figure 5.**
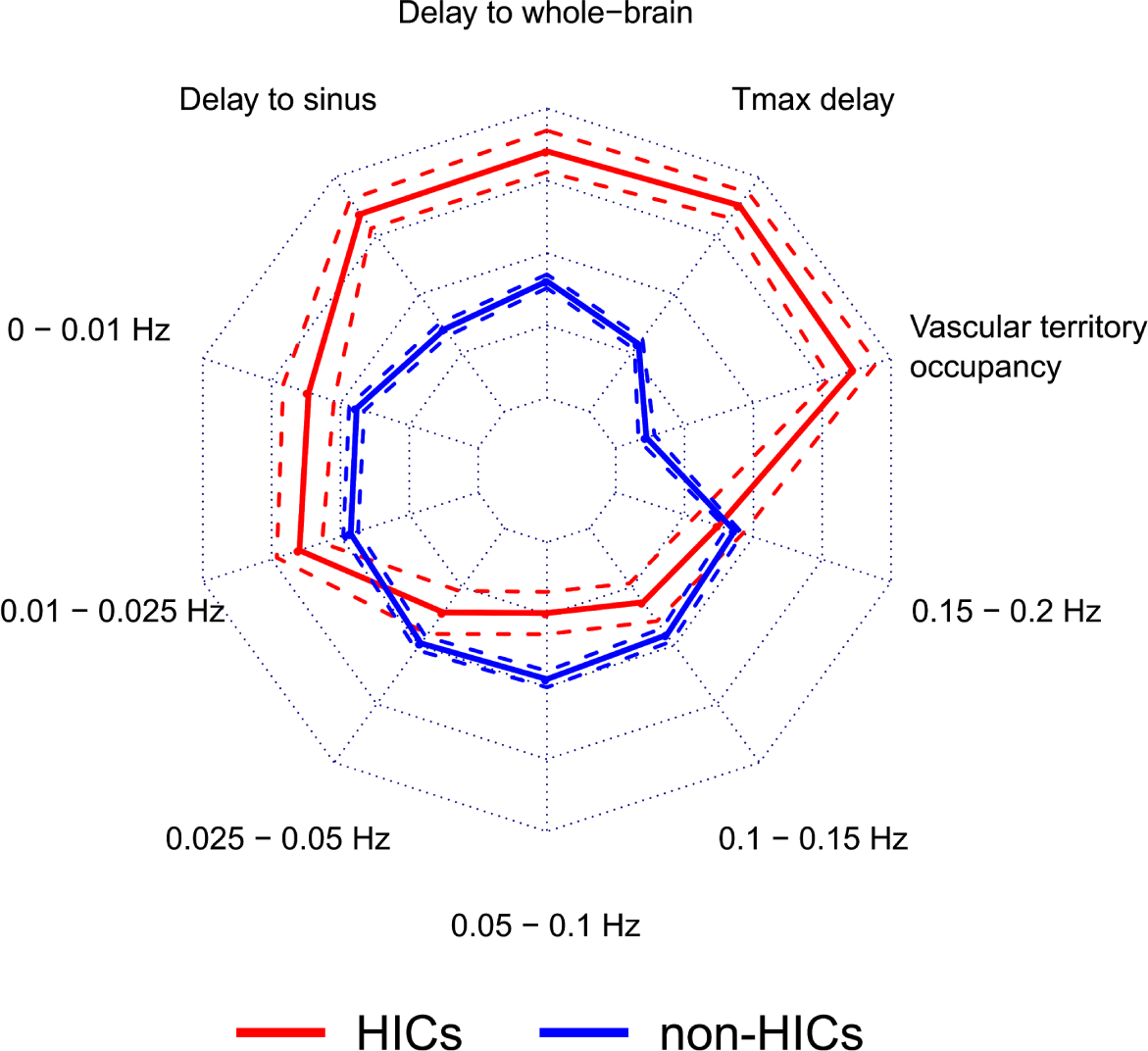
Hypoperfusion independent components (HICs) are characterized by distinct features that distinguish them from non-HICs independent components. This radar plot shows the feature values for HICs (n=57) and non-HICs (n=471). Solid lines represent the mean feature value and dashed lines represent the 95% confidence interval around the mean. Compared to non-HICs within the same patient, HICs have specific characteristics: higher percentage occupancy within a single vascular territory, presence in regions of higher Tmax delay, time courses that show a higher temporal delay to the whole-brain and the venous sinus time courses, and higher signal power in the lowest frequency bands (0 - 0.01 Hz and 0.01 - 0.025 Hz). This combination of features represents a unique signature for HICs.

### 3.3 Hypoperfusion independent components can be automatically identified

Elastic net regularized GLM was applied to the FIX-shortlisted independent components (i.e. the independent components that were classified by FIX as being non-noise) to determine the predictive power of a set of features on HIC classification. The model discriminated HICs from non-HICs with a median accuracy of 0.96, sensitivity of 1.00, and specificity of 0.96. The median Cohen’s kappa for agreement between the model’s classification and the reference classification was 0.78 and the median AUC was 0.95. Figure 6 shows the distribution of these performance metrics across 50 iterations of the model, using different combinations of HICs and non-HICs.

**Figure 6.**
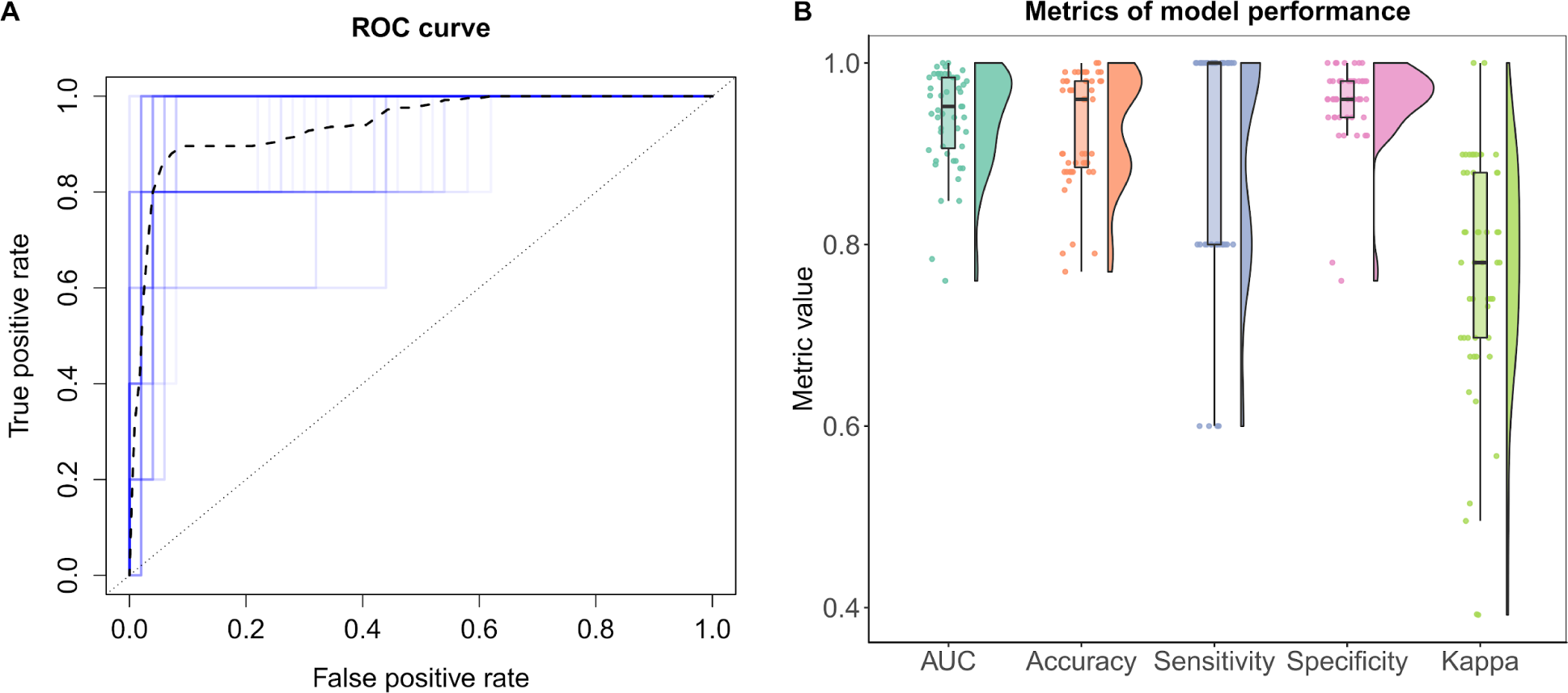
Hypoperfusion independent components (HICs) can be automatically distinguished from other independent components using machine-learning. Using independent test data of different combinations of HICs and non-HICs (that were not used for model training), we ran 50 iterations of the elastic net regularized generalized linear model (GLM) to classify HICs. The overall model performance is shown in the receiver operating characteristic (ROC) curve (A). Each iteration of the model is shown in blue and the mean ROC curve is shown as a black dashed line. The ROC curve depicts the true positive and false positive rates for each classification threshold in the model. The raincloud plot in (B) shows the distribution of different metrics of model performance across the 50 model iterations. The lower and upper hinges of the box plot represent the 25th and 75 percentiles, respectively and the horizontal bar represents the median value. Across all model iterations, the values of area-under-the-ROC-curve (AUC), balanced accuracy, sensitivity, specificity, and Cohen’s kappa are depicted.

A useful feature of elastic net regularized GLM is that it performs implicit variable selection, and thus enhances model interpretability by allowing the assessment of each feature’s relative importance for the classification (Zou & Hastie, 2005). Table 2 shows that 7 out of the 9 extracted features were used by the model to distinguish HICs from non-HICs (i.e. have an Odds ratio ≠ 1). These features included a larger delay in the low frequency oscillations of the HIC relative to the whole-brain and venous sinus signals, a larger restriction to a single vascular territory by the HICs, a higher power in the 0 - 0.01 Hz and 0.01 - 0.025 Hz frequency bands in the HICs, and a higher power in the 0.025 - 0.05 Hz and 0.1 - 0.15 Hz frequency bands in the non-HICs. Note that the implausibly high Odds ratio for the 0.15 - 0.2 Hz variable (2.21 x 10^16^) likely reflects the multicollinearity between the variables describing the power in different frequency ranges (which necessarily exists because these are proportions that sum up to 100%). Importantly, while this means the exact Odds ratios should be interpreted with caution, it does not affect the ability of the model to make predictions (Kutner, MH Nachtsheim, CJ Neter, J Li, W, n.d.).

**Table 2.**
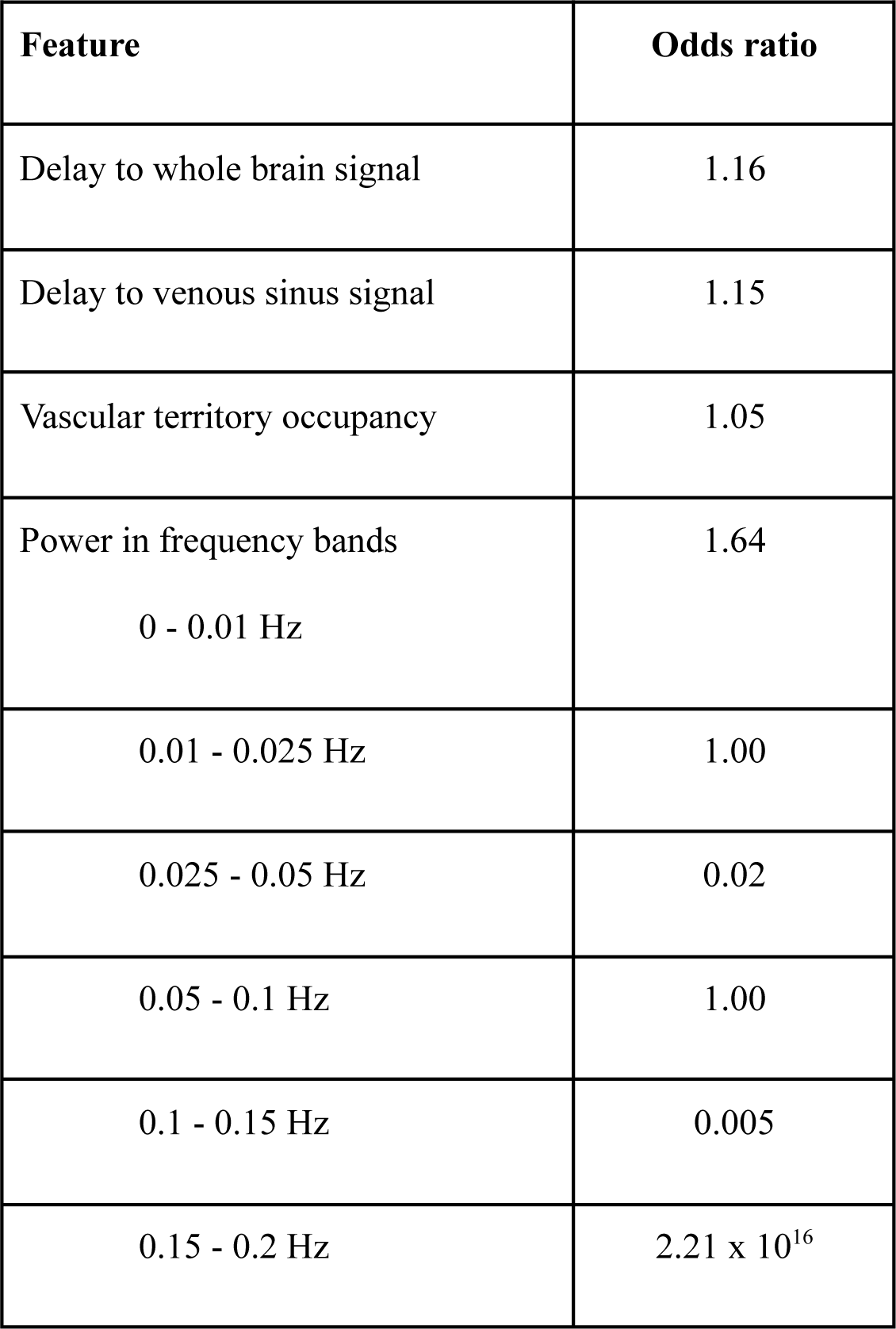
Results of the elastic net regularized generalized linear model for the classification of hypoperfusion independent components (HICs) based on spatial and temporal features.

## 4 Discussion

In this study, we present a novel approach for identifying areas of disturbed perfusion in ischemic stroke without the use of contrast agents. We demonstrate that, using this method, areas of hypoperfusion can be extracted from resting-state fMRI data in a manner that is user-independent and robust to patient motion. The results of this study have important implications from both a clinical and methodological standpoint.

We provide several lines of evidence that certain independent components isolated from rs-fMRI data using spatial ICA reflect regions of hypoperfused brain tissue. Firstly, there is high visual similarity between the spatial maps of HICs and the Tmax maps of individual patients. This is noticeable even when patients had multiple perfusion deficits within (Figure 3) and across (Figure 4) vascular territories. Secondly, HICs were present in areas of much higher Tmax delay than other independent components (Figure 5). Thirdly, HICs observed at baseline disappeared at follow-up in patients with successful recanalization and reperfusion, paralleling the reversal of patients’ DSC-MRI Tmax lesions (Fig 2). Finally, HICs show BOLD signal characteristics that are consistent with those previously described in hypoperfused tissue (Y. Liu et al., 2007; Lv et al., 2013; Tsai et al., 2014; Yao et al., 2012).

These spatial and temporal features of HICs help distinguish them from resting-state networks (Figure 5). BOLD oscillations associated with HICs were more temporally delayed in relation to the global and venous sinus signals than oscillations associated with resting-state networks. Although temporal delays in BOLD LFOs have been shown to occur under physiological conditions (Aso et al., 2017) and differ between resting-state networks (Tong et al., 2015), the largest delays are seen in areas of pathologically reduced perfusion (Khalil et al., 2017, 2020).

Additionally, HICs showed more signal power in very low frequency ranges (0 - 0.025 Hz) and less signal power in the 0.025 - 0.05 Hz range than resting-state networks. This finding is consistent with previous studies showing increased signal power in the <0.01 Hz range in hypoperfused tissue (Y. Liu et al., 2007; Tsai et al., 2014; Yao et al., 2012). Our study adds to this by finding that oscillations associated with resting-state networks exist largely towards the higher end of the low frequency range than oscillations associated with disturbed perfusion.

One disadvantage of spatial ICA is that it can output potentially dozens of components, and browsing through them to visually identify those likely reflecting hypoperfusion can be time-consuming, require expertise, and be subject to bias. We therefore combined the aforementioned component features with the degree of restriction of the component’s spatial map to a single vascular territory and used this set of features to train an algorithm to automatically distinguish HICs from resting-state networks. The algorithm did this with high (>95%) accuracy, sensitivity and specificity, showing that HICs can be automatically extracted from the rest of the components of spatial ICA.

In this study, the spatial agreement between HICs and hypoperfusion is reflected in the fact that the areas covered by the HICs on Tmax maps showed by far the highest Tmax delay (after removal of artifactual Tmax delays in the CSF). Although quantitative metrics of spatial overlap between Tmax perfusion deficits and HICs could have been calculated, this approach would be susceptible to delineation errors or inconsistencies and the limitations inherent to these metrics (such as a high sensitivity to disagreements), and would depend on the Tmax threshold selected. We believe that the high Tmax delay in the areas covered by HICs, as well as the visual similarity between HICs and Tmax maps (which can be interactively viewed here:https://doi.org/10.6084/m9.figshare.13686931), to be more appropriate evidence for their spatial agreement.

Spatial ICA has certain advantages over existing methods for assessing perfusion using rs-fMRI. Instead of assessing the individual characteristics of hypoperfusion-related signals separately, spatial ICA extracts components that reflect a combination of changes in frequency, amplitude, and temporal delay. In addition, we show that spatial ICA is capable of identifying perfusion deficits even in the presence of severe patient motion, where methods based on cross-correlation, such as time shift analysis (Lv et al., 2013), often fail (Figure 3 and https://doi.org/10.6084/m9.figshare.13676779.v1). Finally, unlike time shift analysis, spatial ICA does not require the specification of a reference signal, the choice of which can substantially affect the calculated maps (Christen et al., 2015; Khalil et al., 2017; J. Wu et al., 2017).

This study’s findings have two major implications. From a clinical perspective, it provides a proof-of-concept for a new method of assessing blood flow in acute stroke patients that may be relevant for informing decision-making regarding recanalization and reperfusion therapies. This method is safer, as it does not require the use of exogenous contrast agents, which are particularly problematic in this patient population, who are generally older and have a higher prevalence of chronic kidney disease (Sadowski et al., 2007). The acquisition of the rs-fMRI data, upon which this method is based, is simple and widely available on clinical scanners. Although the rs-fMRI data acquisition took substantially longer than DSC-MRI in this study (about six minutes vs two minutes), recent evidence suggests that extracting perfusion-related information from rs-fMRI can be achieved with much shorter acquisitions (Tanrıtanır et al., 2020). The method is also user-independent, thereby avoiding the subjective assessment of perfusion maps that often leads to inconsistencies between and within experts (Campbell et al., 2010) and allows it to be seamlessly integrated in routine clinical practice. Finally, the robustness of the method to head motion is a substantial practical advantage, as patients scanned during acute illnesses tend to exhibit a lot of motion, which diminishes scan quality and interpretability (Andre et al., 2015). For further clinical validation of this method, detailed comparisons to quantitative perfusion thresholds derived from DSC-MRI and to imaging and clinical outcomes should be made in future studies.

From a methodological perspective, this study underscores the importance of accounting for disturbed perfusion as a potential source of confounding in rs-fMRI studies, particularly in patients with cerebrovascular diseases. Studies investigating functional connectivity in patients with potentially abnormal cerebral perfusion should therefore attempt to disentangle the effects of disturbed perfusion on the BOLD signal (e.g. due to vessel pathology) from components of the BOLD signal that reflect neuronal activity via local neurovascular coupling. So far, suggestions on how to do this have included regressing out the time delays (relative to a reference) from the rs-fMRI data (Erdoğan et al., 2016) and temporally realigning the BOLD signal time courses according to each voxel’s time delay value (Jahanian et al., 2018).

While the spatial distribution of physiological vascular processes overlap with, and are often indistinguishable from, the spatial distribution of resting-state networks (Bright et al., 2018; J. E. Chen et al., 2020; Tong et al., 2015), we show that the influence of disturbed perfusion on the BOLD signal in stroke patients can be readily disentangled from other components of the BOLD signal using spatial ICA. There are two potential explanations for this. The first is that, in stroke, hypoperfusion is spatially restricted to either a vascular territory or part of a vascular territory. Onthe other hand, physiological vascular processes are spatially distributed in a manner similar to resting-state networks (Bright et al., 2018; J. E. Chen et al., 2020; Tong et al., 2015), which may make their separation using spatial ICA less likely. The second possible explanation has to do with the large difference in the temporal BOLD signal characteristics between hypoperfused and normally perfused tissue, which may also facilitate their separation by spatial ICA. In stroke, disturbed perfusion tends to be severe compared to the physiological delays in perfusion across different brain regions, and this leads to relatively large changes in the temporal characteristics of the BOLD signal in hypoperfused regions (Khalil et al., 2017, 2020; Y. Liu et al., 2007; Tsai et al., 2014; Yao et al., 2012).

As the first study to describe this method for assessing perfusion, the study has some limitations. The study sample is relatively small, owing to the fact that it includes an established, albeit relatively invasive, reference standard for assessing perfusion (DSC-MRI) as a comparison. Because of the novelty of the method, we chose to have the two raters perform the ratings together and therefore could not quantify interrater agreement on the identification of HICs in this study. Future studies should test the algorithm we developed on larger cohorts with different MR sequence parameters and more heterogeneous patient cohorts, assess practical aspects of the method, such as the required computing time and power, and investigate the minimum scan length required for the method to deliver reliable results as has recently been done for time shift analysis (Tanrıtanır et al., 2020). Larger cohorts would also allow the investigation of the clinical significance of perfusion assessed using this method, in terms of how it relates to clinical and imaging outcomes and how it potentially influences clinical decision-making in acute stroke patients (Fisher & Albers, 2013). Only two of the four cases in our cohort where spatial ICA could not identify the perfusion deficit could be explained by severe head motion. Therefore, larger studies should investigate in more detail the causes of such false negative cases. Finally, the nature of ICA means that, depending on the properties of the algorithm used, individual signal sources (such as a resting state network or hypoperfused tissue) can be spread across multiple components (Esposito et al., 2002). Therefore, it is important to note that the term “hypoperfusion independent component” is an operationalization and that other independent components might also partially reflect hypoperfused tissue.

In summary, spatial independent component analysis is a novel approach for identifying hypoperfused tissue in ischemic stroke. It does not require the use of exogenous contrast agents, its data can be analyzed without user input, and its results are robust to patient motion. It therefore presents a convenient and promising new alternative to existing perfusion imaging methods in acute stroke.

## Acknowledgments

The authors thank Prof. Dr. med. Andreas Meisel for providing feedback on the manuscript.

## 7 Author contribution statement

J-Y.H.: conceptualization, methodology, software, formal analysis, visualization, data curation, writing - original draft, writing - review & editing

E.K.: methodology, writing - review & editing T.N.: methodology, writing - review & editing

S.O-C.: visualization, writing - original draft, writing - review & editing M.L.: methodology, formal analysis, writing - review & editing

K.V.: project administration, investigation, writing - review & editing D.M.: methodology, writing - review & editing

J.B.F.: resources, project administration, supervision, writing - review & editing A.V.: resources, supervision, writing - review & editing

A.A.K.: conceptualization, methodology, software, formal analysis, visualization, data curation, supervision, writing - original draft, writing - review & editing

## Data availability

The data and code for the training and testing of the model used in this manuscript can be found at https://github.com/ahmedaak/spatial_ICA_stroke.

## Funding

The authors disclosed receipt of the following financial support for the research, authorship, and/or publication of this article: A.A.K. received funding from the NeuroCure Cluster of Excellence and the Berlin Institute of Health Junior Clinician-Scientist Programme. The funding sources played no role in this study’s design, in data collection, in the analysis or interpretation of the data, in writing this article, or in the decision to submit this article for publication.

## Conflict of interest disclosure

J-Y.H., A.A.K., K.V., and J.B.F. are co-inventors of a method for automatically delineating perfusion lesions on perfusion MRI data (European Patent application 17179320.01-1906), distinct from the method described in this study, which is now in the public domain. J.B.F. reports grants from European Union 7th Framework Program and personal fees from Bioclinica, Artemida, Cerevast, Brainomix, BMS, Merck, Eisai, Biogen, Guerbet, and Nicolab outside the submitted work. All other authors declare that they have no conflict of interest.

## Ethics approval statement

Patients provided written informed consent prior to participation, and all procedures were approved by the local ethics committee and were performed according to the Declaration of Helsinki.

