## Supplemental data for "A novel approach for assessing hypoperfusion in stroke using spatial independent component analysis of resting-state fMRI"

### Supplementary Material

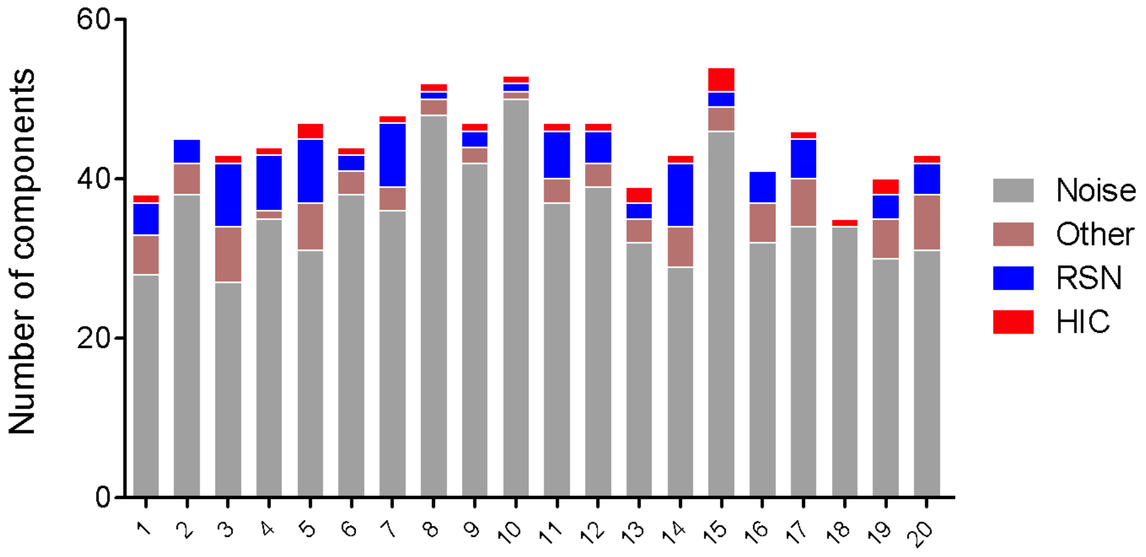


*Supplementary Figure 1.* ***Distribution of independent component classes in the training dataset***. *Bar plot showing the results of the manual classification of independent components for individual patients in the training dataset (n = 20). Each bar represents one rsfMRI scan from one patient. The total number of components differs across patients because MELODIC estimates data dimensionality automatically on a per-subject basis. RSN: resting-state network, HIC: hypoperfusion independent component.*

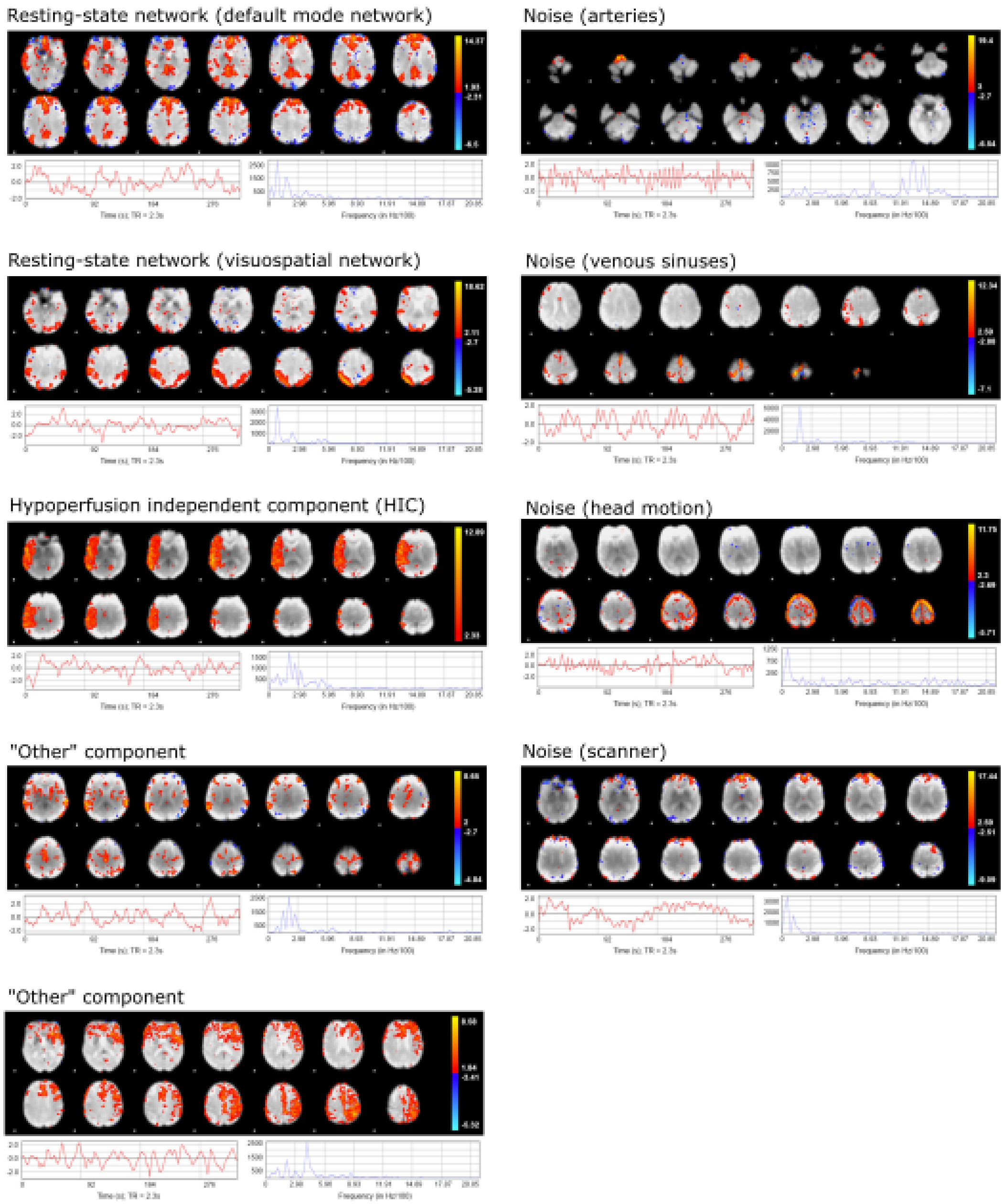


*Supplementary Figure 2.* ***Examples of manual classifications of the output of spatial independent component analysis (spatial ICA)****. Each independent component was classified as either resting state network, hypoperfusion independent component, noise (subdivided into arteries, veins, head motion, or scanner), or other (when unclear). Classification was based on the spatial distribution of the component map, as well as its temporal characteristics.*

*Supplementary Table 1. FMRIB's ICA-based X-noiseifier (FIX) classification accuracies from training data (n=20).*

| **Threshold** | **1** | **2** | **5** | **10** | **20** | **30** | **40** | **50** |
|---|---|---|---|---|---|---|---|---|
| **Mean TPR** | 96.8 | 96.8 | 96.0 | 95.3 | 90.9 | 86.2 | 80.3 | 76.3 |
| **Mean TNR** | 67.3 | 71.1 | 74.9 | 77.8 | 84.6 | 90.1 | 94.0 | 95.6 |
| **Number of subjects with HICs incorrectly classified by FIX as noise** | 0 | 0 | 0 | 0 | 0 | 3 (15%) | 4 (20%) | 4 (20%) |

*TPR: true positive rate; TNR: true negative rate; FIX: FMRIB's ICA-based X-noiseifier. The red box identifies the optimal threshold selected for the classification of independent components in the training dataset.*

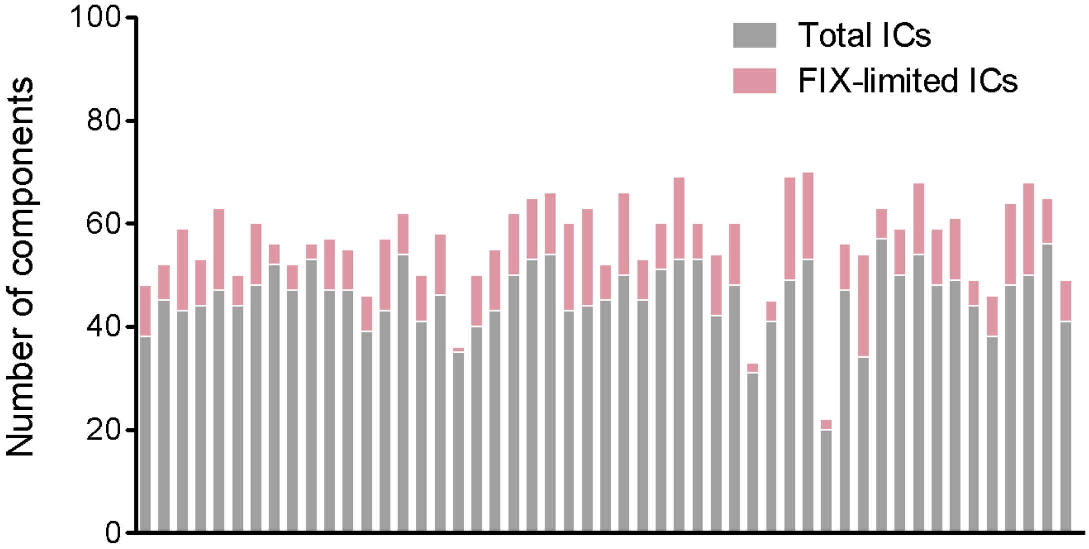


*Supplementary Figure 3.* ***Results of FMRIB's ICA-based X-noiseifier (FIX) denoising of the full dataset.*** *The bar plot shows the number of likely-signal independent components (pink) identified by FIX from the total number of components (grey) for each subject (x-axis) in the full dataset of 51 scans.*

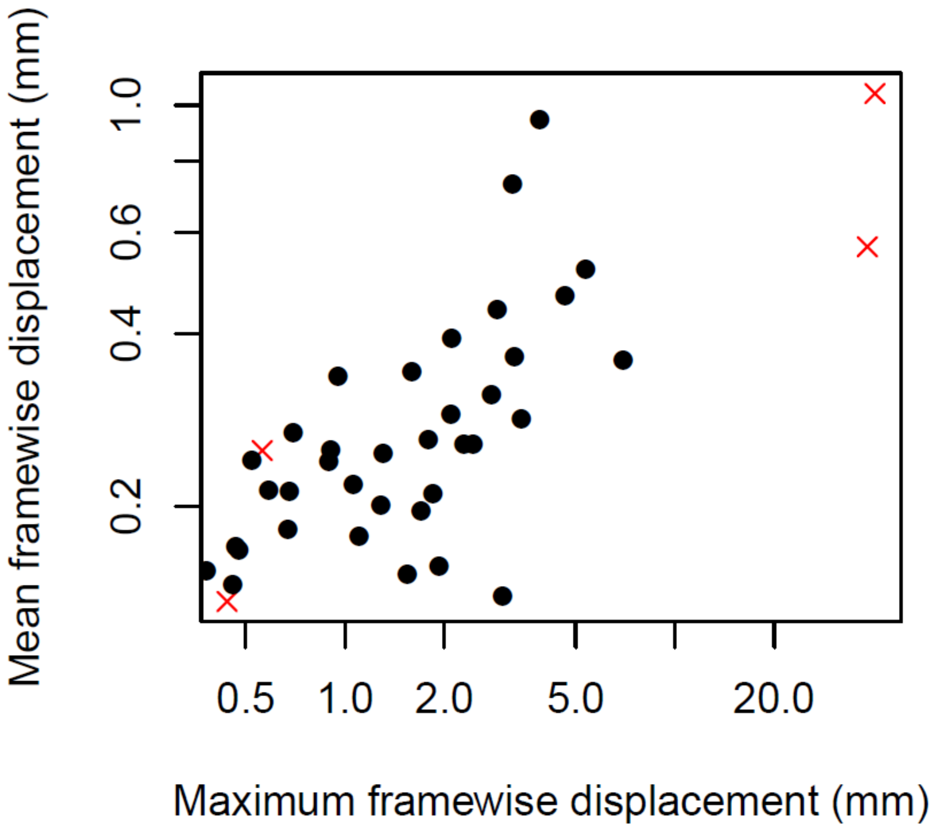


*Supplementary Figure 4.* ***Distribution of head motion during the rs-fMRI scan in the study sample****. Scatter plot showing the distribution of mean and maximum framewise displacement values in the study sample. The data points marked by a red cross correspond to patients in which no hypoperfusion independent component (HIC) was visible despite the presence of a perfusion lesion on Tmax maps (i.e. false negatives). Note that both the x and y axes are on a logarithmic scale.*

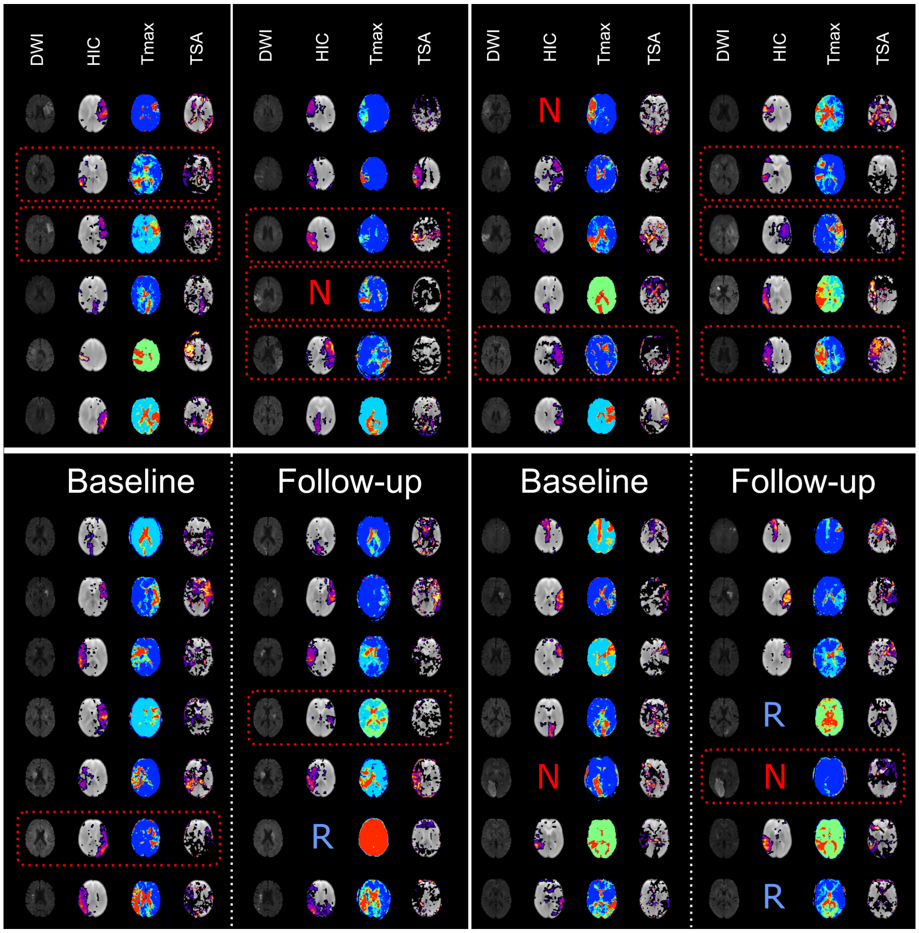


*Supplementary Figure 5.* ***Spatial independent component analysis (spatial ICA) of resting-state fMRI detects post-stroke perfusion deficits in the form of hypoperfusion independent components (HICs).*** *The figure shows diffusion-weighted imaging (visualization of infarcted tissue), time-to-maximum of the residual curve (Tmax; reflecting perfusion), hypoperfusion independent components (HIC), and time shift analysis (TSA) maps from all patients in the study sample (n=37). The top part of the figure shows patients who only received baseline scans (n=23), while the bottom part shows patients who received both baseline and follow-up scans (n=14). An "N" in the HIC panel represents a false-negative; absent HIC despite the presence of a corresponding Tmax lesion. An "R" in the HIC panel (only in the lower part of the figure) represents an absent HIC due to vessel recanalization and tissue reperfusion on the follow-up scans. The dashed red boxes around some patients indicate severe head motion during the rsfMRI scan (> 0.4 mm mean framewise displacement or > 3 mm maximum framewise*

*displacement). Note that, for the time shift analysis maps, only "positive delays" indicating hypoperfused tissue are shown for easier visualization.*
